# Genomic differentiation of the black-chinned tilapia (*Sarotherodon melanotheron*) along a fresh-to-hypersaline water gradient

**DOI:** 10.1101/2025.10.07.680752

**Authors:** Mbaye Tine, Florian Goutieras, Helena D’cotta, Jean-François Baroiller, Simon George, Khalid Belkhir, Jean-Dominique Durand, Catherine Lorin-Nebel, Bruno Guinand

**Author notes:** CHU de Montpellier, Montpellier, France.

## Abstract

The black-chinned tilapia (*Sarotherodon melanotheron*) is an African fish species, found in freshwater, brackish, marine and especially hypersaline (up to 110 ‰) habitats in Senegambia. Using 16,786 filtered single nucleotide polymorphism (SNP) markers, we investigated whether it has responded adaptively to this fresh-to-hypersaline water gradient. Significant genetic differentiation between samples was observed (*F*_ST_ = 0.0568, *p* < 0.01), revealing an interplay between geographic and environmental variation. We focused on a set of outlier SNPs (*n* = 255; *F*_ST_ = 0.320, *p* < 0.001) indicative of adaptive variation, 119 of which mapped to annotated genes in the *Oreochromis niloticus* genome. Significant enrichment was found for physiological pathways relevant to osmosensing and osmoregulation (e.g. inositol phosphate, thyroid hormone synthesis pathways), but also for immune-related pathways that could be activated by ion fluxes (e.g. inflammasome). Some outlier loci mapped to genes that are known to respond to salinity variation in other organisms, including genes found to be differentially expressed in black-chinned tilapia. Adaptive variation along a fresh-to-hypersaline water gradient is well supported in black-chinned tilapia, but its association with climate change specifically induced by hypersalinity deserves further attention particularly in the context of increasing cases of inverted estuaries being reported worldwide.

## 1. Introduction

Salinity plays a central role in shaping the physical (e.g. stratification and dissolved oxygen) and biological (e.g. osmoregulation, thermotolerance, metabolic rates and immunity) environments of estuarine and coastal organisms (Kültz, 2015; Röthig et al., 2023). Changes in salinity are recognised as a key ecosystem process that is affected by climate warming (Chilton et al., 2021; Gillanders et al., 2022; Bueno et al., 2025). Consequently, salinity increases in estuaries are expected to become more widespread as a result of global climate change (Lee et al., 2025). Increases in salinity are already commonly reported in subtropical and Mediterranean estuaries due to a negative water balance caused by increased evaporation and reduced precipitation. However, this is also influenced by sea level rise and changes in tidal amplitude (Hallett et al., 2018; Tweedley et al., 2019). This includes West African estuaries and coastal areas (Tweedley et al., 2019), where, in the most extreme cases, the classic freshwater–seawater (FW–SW) salinity gradient is reversed between upstream reaches and the estuary. This leads to the so-called ‘inverse estuary’, where hypersaline conditions (i.e., >40 ‰) develop in the upper reaches (Potter et al., 2010). Research on estuarine organisms has shown that hypersalinity has a detrimental effect on organism richness, abundance, biomass, community composition, functional diversity, and ultimately, ecosystem health (Forbes & Cyrus, 1993; Kantoussan et al., 2012; Veale et al., 2014; Villanueva, 2015; Breaux et al., 2019; Shadrin & Anufrieva, 2020; Getz & Eckert, 2023; Miosley et al., 2023). Nevertheless, hypersaline systems could be considered natural laboratories providing opportunities to study how organisms cross intensity thresholds inducing energetic constraints and physiological stress and successfully adapt to stressful aquatic environments (Saccò et al., 2021; Esbaugh, 2025).

Advances in genomics are providing access to markers that help us to better understand how organisms cope with extreme environments induced by climate change (Wang & Guo, 2019; Bernatchez et al., 2024). Thus, improved knowledge of genome-wide molecular responses may provide valuable insights into potential adaptations across an extended salinity gradient, which exerts strong selective pressure on traits linked to osmoregulation and ionoregulation. However, most studies investigating the response to salinity challenges using transcriptomic or genomic tools have focused on the FW-SW gradient (Velotta et al., 2022; Ahi et al., 2025 for reviews). In contrast, few studies have examined the response to hypersalinity using high-throughput proteomics or transcriptomics (Lam et al., 2014; Schauer et al., 2018; Su et al., 2020; Root & Kültz, 2024; Tao & Breves, 2024), and they rarely considered wild samples (Li and Kültz, 2020; Blondeau-Bidet et al., 2024; Wilson et al., 2024). A genomic signature of adaptive variation at the DNA level, associated with a freshwater-hypersaline gradient, could illustrate impending physiological threats associated with climate change in fish.

The black-chinned (or blackchin) tilapia *Sarotherodon melanotheron* (Rüppell, 1852) is a euryhaline, mouth-brooding species found from southern Cameroon to Senegal. It is widely exploited in artisanal fisheries that support local livelihoods and food security (e.g. Lederoun et al., 2020; Hountcheme et al., 2025). It is naturally distributed in freshwater (FW), coastal seawater (SW) and brackish environments, and is notable for its ability to tolerate and acclimatise to hypersaline conditions in some estuaries and lagoons (Panfili et al., 2004, 2006; Amoussou et al., 2018). This is evident in Senegambia. In this area, the Saloum estuary — a drowned river valley — and the Casamance River became permanent inverted estuaries. The first signs of increasing salinity in the Saloum date back to the second half of the nineteenth century (Carré et al., 2019), but hypersalinity mainly occurred after Sahelian droughts affected West Africa in the late 1960s and early 1970s (Pagès & Citeau, 1990; Savenije & Pagès, 1992). In the Saloum estuary, the salinity of the landward waters can be up to three times higher than that of seawater during the dry season, and rarely falls below 40 ‰ during the rainy season (Panfili et al., 2004, 2006; Descroix et al., 2020). The black-chinned tilapia is the only fish species that inhabits the upper reaches of the Saloum estuary permanently during the dry season, when salinities are at their highest (Panfili et al., 2004), and it can reproduce at salinities of up to 60–70 ‰ (Guèye et al., 2012). A similar, albeit less pronounced, gradient (up to 70 ‰) is observed in the Casamance River (Debenay et al., 1994; Labonne et al., 2009). By contrast, neighbouring Senegambian estuaries where the black-chinned tilapia is also present function normally (e.g. the Gambia and Senegal rivers), with salinity increasing from the upper to the lower reaches due to increased FW discharge or dams limiting seawater intrusion (Gac et al., 1986; Descroix et al., 2020). A large number of field and laboratory studies of black-chinned tilapia from this area have demonstrated phenotypic changes in response to increased salinity. These include osmoregulatory performance (Ouattara et al., 2009; Lorin-Nebel et al., 2012; Riou et al., 2012), sperm motility and activation (Legendre et al., 2016), as well as condition, life history, growth rate, and reproductive traits (Diop et al., 2004; Panfili et al., 2004, 2006; Diouf et al., 2009; Labonne et al., 2009; Guèye et al., 2012, 2016, 2020; Dugué et al., 2014). Salinity-induced phenotypic responses have also been studied at the molecular level, primarily focusing on the differential expression of candidate genes in various organs, such as the gills (Tine et al., 2007, 2008, 2010, 2011, 2012), the intestine and the liver (Link et al., 2010), the gonads (Avarre et al., 2014a, b; Link et al., 2022) and the immune organs (Link et al., 2022). Recently, a high-throughput RNA sequencing approach was used to analyse the gills of *S. melanotheron*, revealing genome-wide changes in gene expression between wild populations distributed over a salinity range of 0–80 ‰. The study also investigated how canonical biological pathways respond to hypo- and hyperosmotic stress (Blondeau-Bidet et al., 2024). Taken together, these studies suggest that local adaptation across an extended salinity gradient may occur in Senegambian populations of black-chinned tilapia. However, the observed changes may simply reflect phenotypic plasticity (i.e. physiological flexibility) or gene-environment interactions. Salt tolerance in tilapia is indeed a complex trait influenced by a combination of genetic components and environmental conditions (Yue et al., 2024). Furthermore, the presence of a genome-wide signature of salinity-based adaptation (i.e. isolation by environment, whereby genetic differentiation increases with environmental differences, regardless of geographic distance; Wang & Bradburd, 2014) has yet to be adequately investigated in black-chinned tilapia. Agnèse et al. (2008) reported possible adaptive variation at a microsatellite locus within the promoter of the ‘FW adaptation hormone’ prolactin (Manzon, 2002), which is associated with salinity conditions in a landlocked population of *S. melanotheron* in the Ivory Coast. However, this study did not consider hypersaline conditions. No genome-wide study based on DNA markers has yet been carried out in this species.

This study aims to investigate neutral and adaptive genetic variation at single nucleotide polymorphisms (SNPs) across the genome using restriction site-associated DNA sequencing (RADseq). Six samples of black-chinned tilapia from Senegal and The Gambia, covering freshwater, marine coastal and hypersaline estuarine habitats, will be analysed. This will enable us to gain insight into the potential contribution of salinity adaptation along a fresh-to-hypersaline water gradient to the genetic differentiation of this species within West African estuarine ecosystems impacted by climate change.

## 2. Materials and Methods

### 2.1. Sampling

Individuals of *S. melanotheron* (*N* = 84) were sampled at the end of the dry season in 2006, with size (fork length) varying between 120 and 160 mm (Tine et al., 2011). Individuals belong to the subspecies *S. m. heudelotii* (Duméril, 1859), which occurs in Senegambia (Falk et al., 2003). Unless otherwise stated, we used *S. melanotheron* to refer to this subspecies. Individuals originated from six localities distributed at different sites in Senegal (Saloum estuary, Lake Guiers) and the Gambia (River Gambia), with one fully marine coastal population (Hann Bay, Dakar) (Fig. 1). These locations are characterised by distinct salinity levels and salinity variations throughout the year in their respective environments (Fig. 1; Tine et al., 2011). Briefly, Lake Guiers is entirely FW, while Hann Bay is a permanent SW environment. Both show no spatial or temporal salinity variation. The remaining sites in Gambia and the Saloum are subject to seasonal variations in salinity, with dry and rainy conditions of the monsoon regime: Balingho (Gambia) is a freshwater location that experiences brackish conditions during the dry season (range: 0-25 ‰), while the Saloum classically supports salinities equal to seawater or hypersalinity during the dry season, and brackish conditions at upstream reaches during the rainy season due to rainfall (minimum salinity: 20-25‰; Panfili et al., 2006). Freshwater conditions are not encountered in the Saloum. Similar ranges for salinity have been reported in other studies in the same area for other sampling years (Savenije et al., 1992; Albaret et al., 2004; Panfili et al., 2004; Simier et al., 2019; Descroix et al., 2020). A description of basic environmental parameters (salinity and temperature in both the dry and rainy seasons) at time of sampling (Tine et al., 2011) is reported in Suppl. Mat. Table S1. Tissue sample preservation is also described in Tine et al. (2011).

**Fig. 1:**
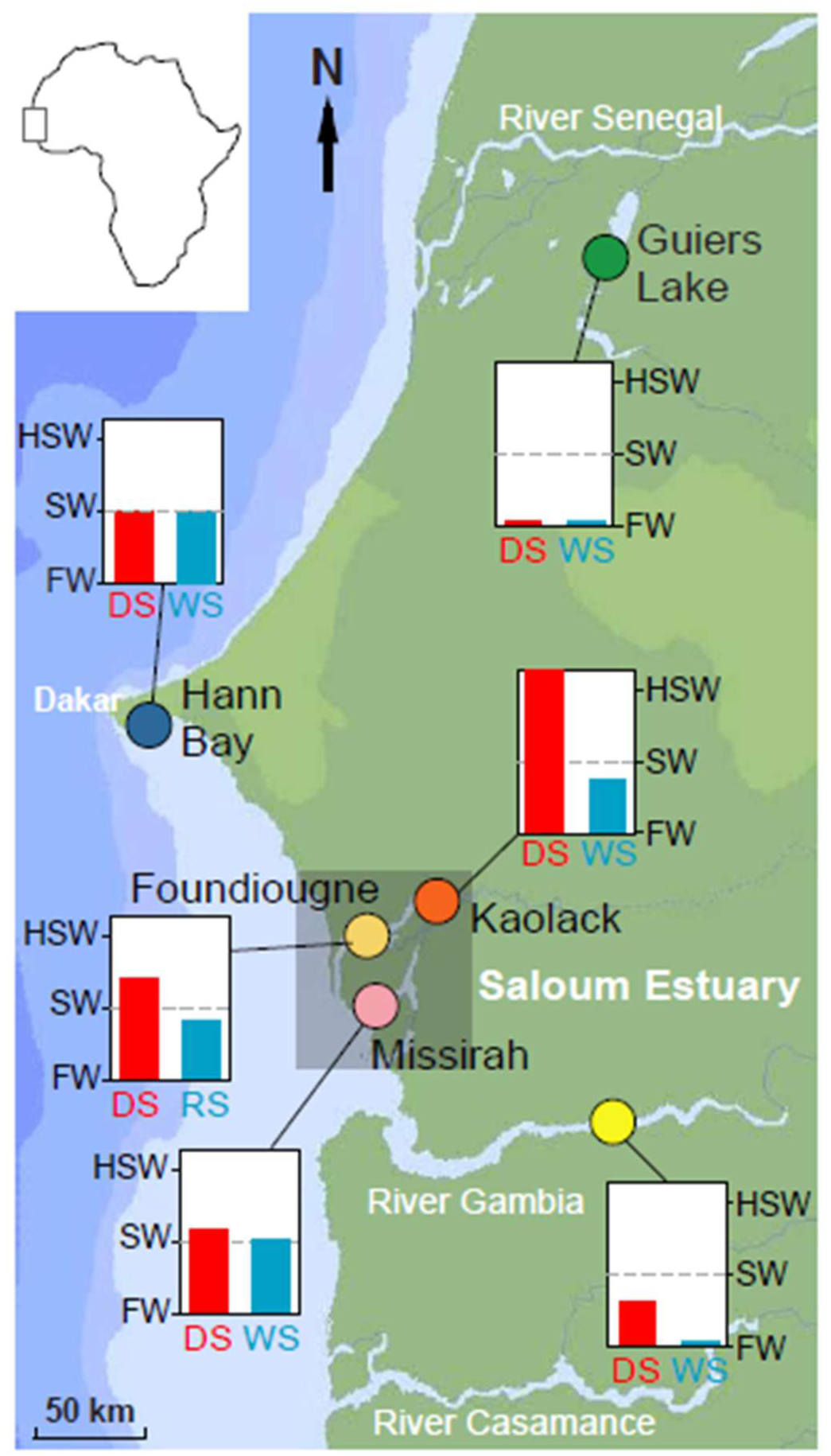
Localization and map of the sampling area in Senegambia. A dark grey shadow indicates samples of the inverse Saloum Estuary. A diagrammatic representation of the salinity variation between the dry and rainy season (DS and RS, respectively) is given for each sample and approximately illustrate conditions encountered at time of sampling (2006). FW: Fresh water; SW: seawater; HSW; hypersaline water.

### 2.2. DNA extraction and library construction

Gill tissues stored at -80°C since 2006 were used for genomic DNA (gDNA) extraction. gDNA was isolated on a KingFisher Flex robot (ThermoFisher, USA) using the Nucleomag 96 Tissue Kit (Macherey-Nagel, Germany) according to the manufacturer’s instructions. DNA quality was assessed by agarose gel electrophoresis and quantified using the dsDNA BR Qubit Quant-iT assay (Invitrogen, France). A single RADseq library was prepared at Montpellier Genomix (MGX, University of Montpellier, France) using Illumina Nextera XT barcodes according to the protocol described in Baird et al. (2008). Briefly, an equal amount of 1 μg DNA from each individual was digested with the enzyme SbfI HT (New England Biolabs), followed by sequencing adapter ligation. Modified Illumina adapters containing five nucleotides of barcode sequence (P1 adapters) unique to an individual in the library were ligated with T4 DNA ligase (New England Biolabs) to multiplex samples. The ligated DNA samples were pooled at an equimolar concentration and sheared to an average size of 1000 base pairs using an ultrasonicator (Covaris). Additional size selection of the pooled DNA was performed by automated sizing (Pippin HT) on an agarose gel. The NEBNext ultra II DNA library preparation kit (New England Biolabs) was used for end repair, A-tailing and ligation of P2 adapters. Reactions were purified using AmpureXP beads (Beckmann Coulter). Samples were amplified using Phusion High-Fidelity PCR Master Mix by 12 cycles of PCR and the library was finally purified using a MinElute column (Qiagen) to obtain approximately 20µL of sequencing library. This library was sequenced on a single lane of Illumina NovaSeq 6000 using single end sequencing (100bp).

### 2.3. Data processing

Initial quality control of raw RAD sequencing data was performed using FastQC (Barbraham BioInformatics; www.bioinformatics.babraham.ac.uk). The raw reads were processed using Stacks v2.55 (Catchen et al., 2013) for demultiplexing based on their barcodes, and trimmed to 95 base pairs using the ‘process_rad_tags’ script. The trimmed reads were deposited in the National Center for Biotechnology Information (NCBI) BioProject database under SRA accession number PRJNA1143382. As no genome is available for *S. melanotheron*, but a similarity in the number of chromosomes/linkage groups (LG) is recognized across cichlids (*n* = 22; Conte et al., 2019), the cleaned reads were aligned to the Nile tilapia (*Oreochromis niloticus*) reference genome (accession number: GCF_001858045.3) using Bowtie (Langmead et al., 2009). The BAM files were sorted, indexed and filtered for redundant reads using SAMtools (Li et al., 2009) to ensure accurate detection results. SNP variation detection was then performed using ‘Gstacks’, and the ‘populations’ module of Stacks was used to filter loci (Catchen et al., 2013). Retained filters included *(i)* a minimum percentage of 80% of genotyped individuals in a population to process a locus for that population, *(ii)* the locus must be present in all samples, *(iii)* a maximum heterozygosity of 0.6 to limit impacts of paralogues, and *(iv)* a minimum minor allele frequency of 5% in at least one sample. To account for linkage disequilibrium between SNPs, only one SNP per RAD locus was retained. Variants were collected and filtered using VCFtools (Danecek et al., 2011) for additional SNP quality control. SNPs were then filtered according to sequencing depth, missing data, and number of alleles per site. We only considered sites with depth ≥ 6 across the entire data set, quality score ≥ 30, maximum missing ≤ 0.5, and only 2 alleles present.

### 2.4. Statistical analysis

We computed the inbreeding index *F*_IS_ for each sampling location using the *radiator v1.1.7* package (Gosselin, 2017; *summary_rad* functions). This package was also used to convert the VCF to the *genepop* format used in the package *genepop v1.2.2* (Rousset et al., 2020) in which deviations of Hardy-Weinberg expectations (HWE) were estimated with the *test_HW* function for each locus within each population. SNPs that did not conform to HWE in all six samples were discarded prior to compute overall deviations of HWE for each population by combining results across remaining SNPs using Fisher’s method. Locus specific *F*_ST_ values for all pairwise population comparisons and the weighted average *F*_ST_ values (Weir & Cockerham, 1984) between all population pairs were calculated in ARLEQUIN (Excoffier & Lischer, 2010) after conversion of the format of input files using PGDSPIDER2.0 (Lischer & Excoffier, 2012). Significant testing was assessed with 10,000 permutations. Isolation-by-distance signals were tested using a correlation between *F*_ST_/(1-*F*_ST_) and log(geographic distance) (Rousset, 1997) in *genepop v1.2.2*. An approximate linear geographic distance following coastlines, estuaries or rivers was retained. A neighbor-joining (NJ) tree was built to cluster individuals according to their respective similarity using the ‘ape’ package (Paradis et al., 2004). Admixture was run over *K* values of three through six, with 10-fold cross validation and 1,000 bootstrap replicates for parameter standard errors (Alexander et al., 2009). We performed a principal component analysis (PCA) to describe population structure, then we detected outlier SNPs that were more differentiated than under a neutral model (Luu et al., 2017). The PCADAPT package v4.3.3 (Privé et al., 2020) was used to clean the data set for the number of loci involved in downstream analysis and to retain the most appropriate number of clusters in the scree plot, which displays in decreasing order the percentage of variance explained by each principal component. However, we did not follow classical rules and settings of PCAADAPT and we classified as outliers those SNPs when they loaded at ± 3 standard deviations (SD) from the mean as traditionally suggested for other analysis (e.g. redundancy analysis; Capblancq et al., 2018). This ±3 SD rule undoubtedly biased the selection of loci with high *F*_ST_ values due to divergent selection (adaptive variation) and neglected processes implying weak or background selection, but it is reasonable for an exploratory analysis.

Genes that were shown to contain putative outlier SNPs were tested for functional enrichment using Metascape v3.5 (Zhou et al., 2019) (https://metascape.org/gp/index.html#/main/step1). The human genome was used as a reference. Metascape refers to different ontologies to detect statistically enriched terms including, e.g., the Gene Ontology (GO), Kyoto Encyclopedia of Genes and Genomes (KEGG) (hsa), Reactome Pathway (R_hsa for *H. sapiens*) and Wiki Pathways (WP) databases. Significantly enriched terms/pathways were identified using a false discovery rate (FDR) approach. In Metascape, all statistically enriched terms were first identified across the same databases, but cumulative hypergeometric *p*-values were calculated and used for filtering. Remaining significant terms were then hierarchically clustered based on kappa statistical similarities (Cohen, 1960). Within each cluster, the enriched term with the highest significance (i.e., highest *–log_10_(P)*) has its generic name retained by Metascape v3.5 to describe this cluster and the genes it contains. Other terms and names might be sometimes more indicative than this generic name and were also inspected.

## 3. Results

We obtained 410,146,622 total sequencing reads of which 69.5% (285,050,067) were retrieved after trimming of the library. After demultiplexing, read numbers per sample ranged from 365,694 to 5,291,007. Three individuals over 84 (3.57%; one individual from Missirah, two individuals from the Lake Guiers) with less than 600,000 reads per individual were discarded from subsequent analyses (Suppl. Mat. Table S1). Individual call rates of the remaining individual were all >95% (Suppl. Mat. Table S1).

### 3.1. Genetic population structure

We first obtained 17,365 SNPs at independent RAD locus that were filtered to 16,786 distinct SNPs after removing markers that did not conform to HWE in all six samples and to PCAADAPT. These SNPs reported a mean genetic population differentiation estimated to *F*_ST_ = 0.0568 (*p* < 0.01) among samples and a distribution of *F*_ST_ values ranging from 0.000 to 0.904 at individual SNPs (Suppl. Mat. Fig. S1). Estimates of *F*_IS_ showed significant departures from HWE in four out of six samples (Table 1).

**Table 1:** Pairwise *F*_ST_ values among black-chinned tilapia samples using (A) the full and (B) neutral (i.e. not containing outliers SNPs) data sets considered in this study. Comparisons among Saloum samples are given in grey. *F*_IS_ values are also given and significance illustrates departures of Hardy-Weinberg expectations.

| **A - All loci** | BA | MI | FO | KA | HB | GL |
| --- | --- | --- | --- | --- | --- | --- |
| BA |  | **0.0367** | **0.0380** | **0.0294** | **0.0622** | **0.0356** |
| MI |  |  | 0.0127 | 0.0164* | **0.0570** | **0.0408** |
| FO |  |  |  | 0.0107 | **0.0535** | **0.0427** |
| KA |  |  |  |  | **0.0553** | **0.0385** |
| HB |  |  |  |  |  | **0.0673** |
| $F_{IS}$ | -0.022 | **0.072** | **0.085** | **0.101** | **0.090** | 0.062 |
| **B - Neutral set** | BA | MI | FO | KA | HB | GL |
| BA |  | **0.0267** | **0.0286** | **0.0265** | **0.0432** | **0.0340** |
| MI |  |  | 0.0106 | 0.0123 | **0.0367** | **0.0363** |
| FO |  |  |  | 0.0085 | **0.0351** | **0.0371** |
| KA |  |  |  |  | **0.0405** | **0.0368** |
| HB |  |  |  |  |  | **0.0497** |
| $F_{IS}$ | 0.020 | **0.076** | **0.077** | **0.096** | **0.103** | 0.054 |

Significant values ( $P < 0.05$ ) are given in bold
\*: marginally significant ( $P = 0.081$ )

The pairwise *F*_ST_ values (Table 1) and the NJ tree (Fig. 2A) revealed that individuals mostly grouped per sample and basin of origin, with Hann Bay being the most significantly differentiated from other samples (Fig. 2, Fig. 3A). The Admixture analysis largely confirmed this observation suggesting the likely existence of *K* = 4 clusters in our samples (Fig. 2B). As the southernmost (Balingho) and northernmost (Lake Guiers) populations grouped together and presented a low pairwise *F*_ST_ value; no pattern of isolation by distance was detected among samples (*r*^2^ = 0.140, *p* = 0.362). The PCA also confirmed these findings, with PCA1 and PCA2 explaining 4.7% and 2.5% of overall genetic variation, respectively (Fig. 3A). Nevertheless, *(i)* PCA2 remarkably showed that samples were broadly ordinated along a fresh-to hypersaline water gradient (Fig. 3A), and, contrary to former analyses, *(ii)* PCA3 that explained 2.0% of overall genetic variation allowed to more clearly illustrate the genetic differentiation between the Balingho (in yellow) and Lake Guiers (in green) samples (Fig. 3B).

**Fig. 2:**
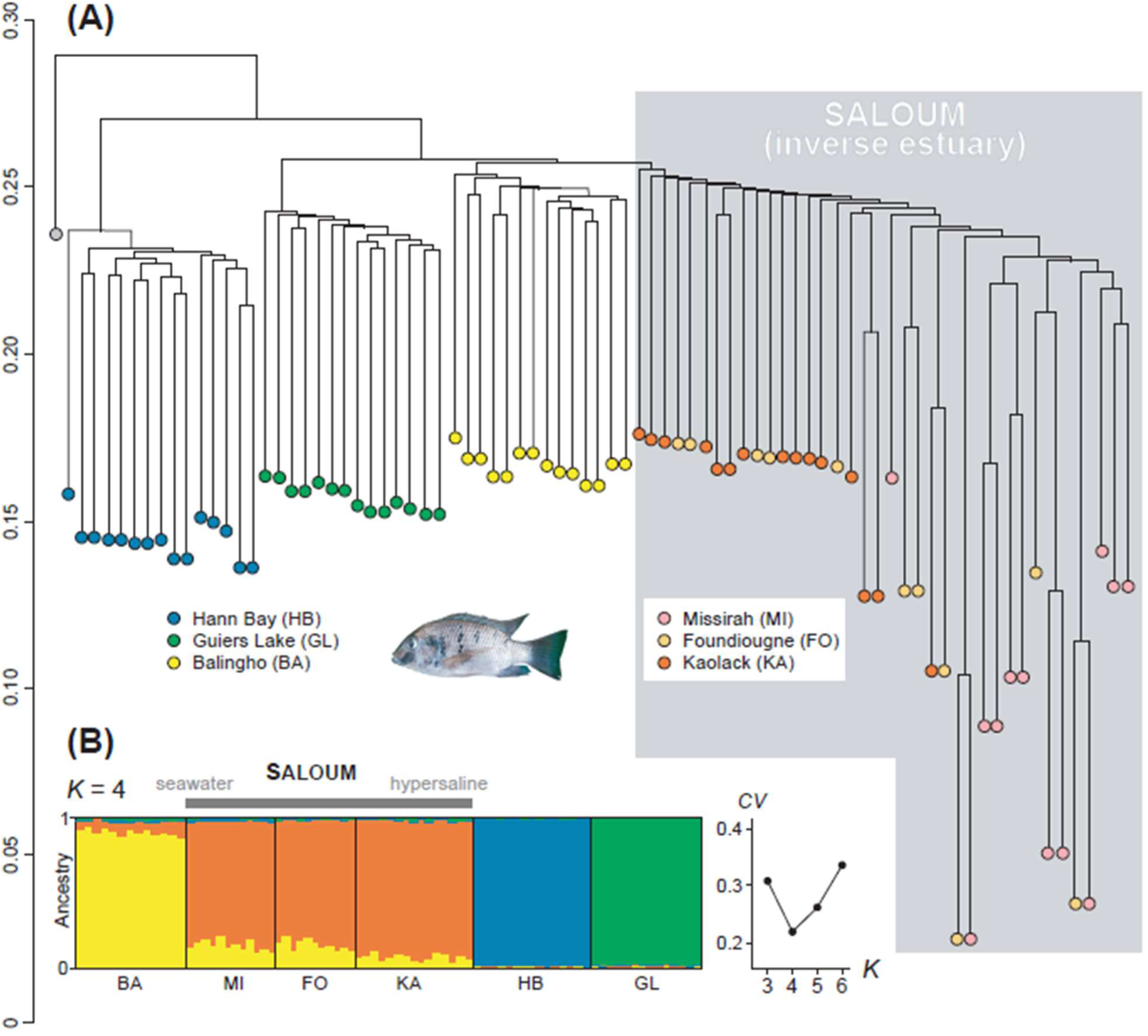
(A) Neighbor-joining tree of the 81 samples retained in this study. The tree is rooted with an individual that has been discarded. (B) Individual ancestry estimated by Admixture for *K* = 4 for which the cross-validation criterion was minimized. BA: Balingho, MI: Missirah, FO: Foundiougne, KA: Kaolack, HB: Hann Bay, GL: Guiers Lake. Color codes as in Fig. 1.

**Fig 3:**
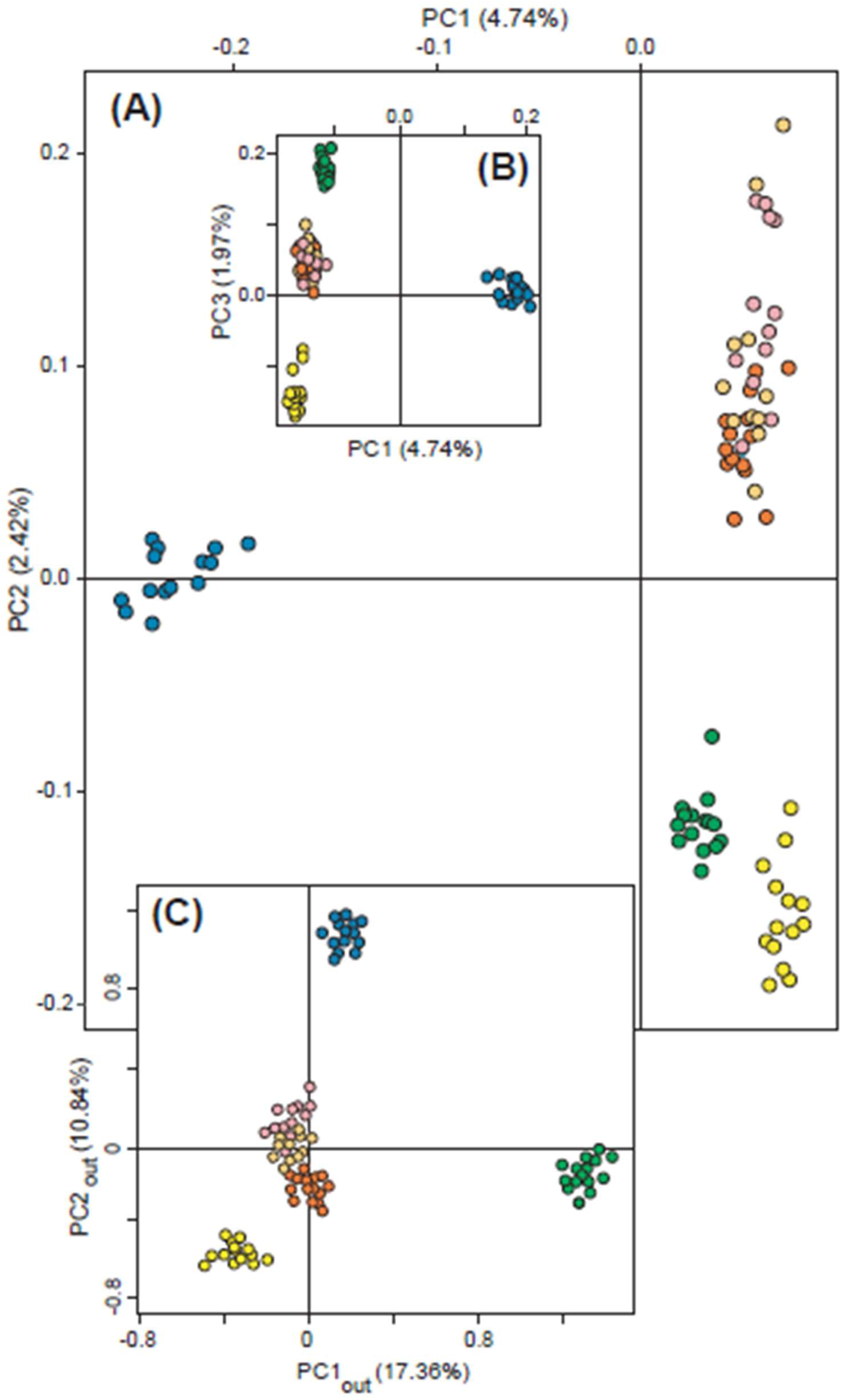
PCA scatterplots – (A) Ordination of individual samples on PC axes 1 and 2 and (B) PC axes 2 and 3 based on the 16,786 SNPs retained in this study. (C) Ordination of samples based on the 255 outlier loci identified in this study. Color codes as in Fig. 1.

### 3.2. Detection of outlier loci

Two hundred and fifty-five outlier loci with ±3 SD deviations from the mean were identified with the PCA approach (1.52% of the total number of SNPs). Outlier loci were found to be distributed over all Nile tilapia LGs and some unassembled scaffolds (Suppl. Mat. Table S2), but numbers of outlier SNPs per LG was not found to be correlated with LG length (*r*^2^ = 0.010, NS). Mean *F*_ST_ value of PCA-based outlier loci was found equal to 0.320 (*p* < 0.001). A PCA conducted on the set of outlier loci is given in Fig. 3C. Results opposed FW Lake Guiers and SW Hann Bay samples to southern ones, with Saloum samples found close but distinct from Gambia samples (Fig. 3C). Among PCA-based outlier loci, twenty-one were found to map to long non-coding RNAs (lncRNAs) (*n* = 16), pseudogenes (*n* = 4) or microRNAs (*n* = 1), while 136 mapped to loci that referred to 119 distinct annotated genes in the Nile tilapia genome (Suppl. Mat. Table S2). Because PCA-based outlier loci are indicative of divergent selection and affect the estimation of population structure indices, we recomputed *F*_IS_ and (pairwise) *F*_ST_ values after having deleted these loci. Original levels of significance using the full set of loci were not modified by the removal of outlier loci (corrected mean *F*_ST_ = 0.0480; *p* = 0.011 for the putative neutral loci), neither for *F*_IS_ nor for pairwise *F*_ST_ (Table 1). The overall pattern of isolation-by-distance across populations remained non-significant once outlier loci were removed from the data set (*r*^2^ = 0.188, *p* = 0.517).

### 3.3. PCA-based enrichment analysis

The list containing the 119 gene names identified with PCA was used in Metascape for functional enrichment analysis. Table 1 shows the top 20 significantly enriched functional terms, some of which are relevant to processes related to the organism’s response to environmental change, but also linked to immunity and development (Fig. 4). Relevant pathways mainly included the two pathways associated with the regulation of assembly/disassembly of proteins (GO:0043254; GO:0043243), protein modification by small protein conjugation or removal (GO: 0070647), the inositol phosphate metabolic process (GO:0043637), but also phospholipase D signaling (R_hsa4072), thyroid hormone synthesis (hsa04918), cell substrate adhesion (GO:0031589), hemostasis (hsa109582), endopeptidase activity (GO:0052548), and the mitotic life cycle (GO:0010389) (Fig. 4). Enriched immune functions were found to refer to the negative regulation of NF-κB (nuclear factor-kappa B) signal transduction pathway (GO:1901223), the regulation of the NLRP3 inflammasome complex assembly (GO:1900225), the endoplasmic reticulum stress response to coronavirus infection (WP4861), and the negative regulation of viral process (GO:0048525) (Fig. 3). Details concerning high *F_ST_* outlier-containing genes involved in each of these functional terms have been reported in the PCA-based Metascape analysis output in Suppl. Mat. Table S3.

**Fig. 4:**
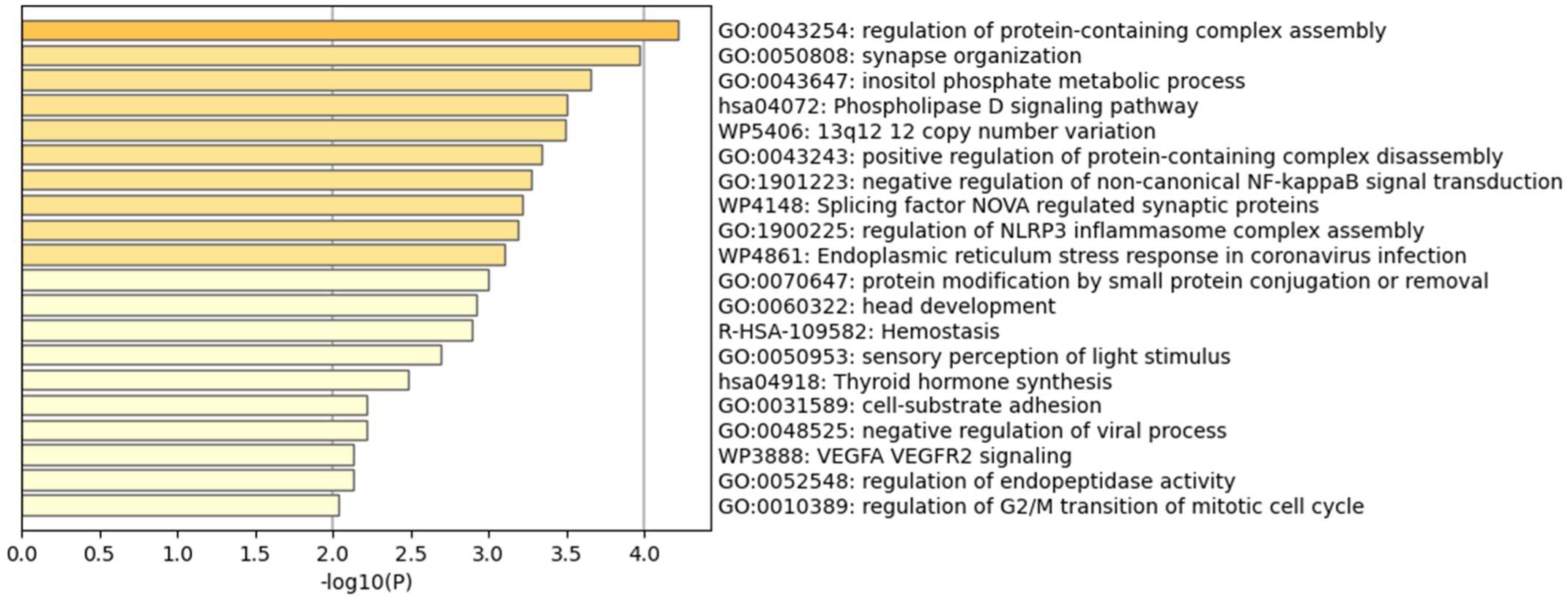
Top 20 significantly enriched pathways based on the 119 annotated genes containing outlier SNPs detected with Metascape. Details are given in Supplementary Table S3.

Based on literature search, the relevance of some mapped outlier-containing genes found in this study is given in Table 2. These genes have been found to be significantly differentially expressed, to contain outlier SNPs and sometimes to have alternatively spliced variants in euryhaline aquatic species (mostly fish), but also, in a few cases, in cyanobacteria and terrestrial plants subjected to salt stress that are involved in mitochondrial homeostasis associated with regulation of free amino-acid osmolytes (*SHMT2*, *PFN1*; Table 2).

**Table 2:**
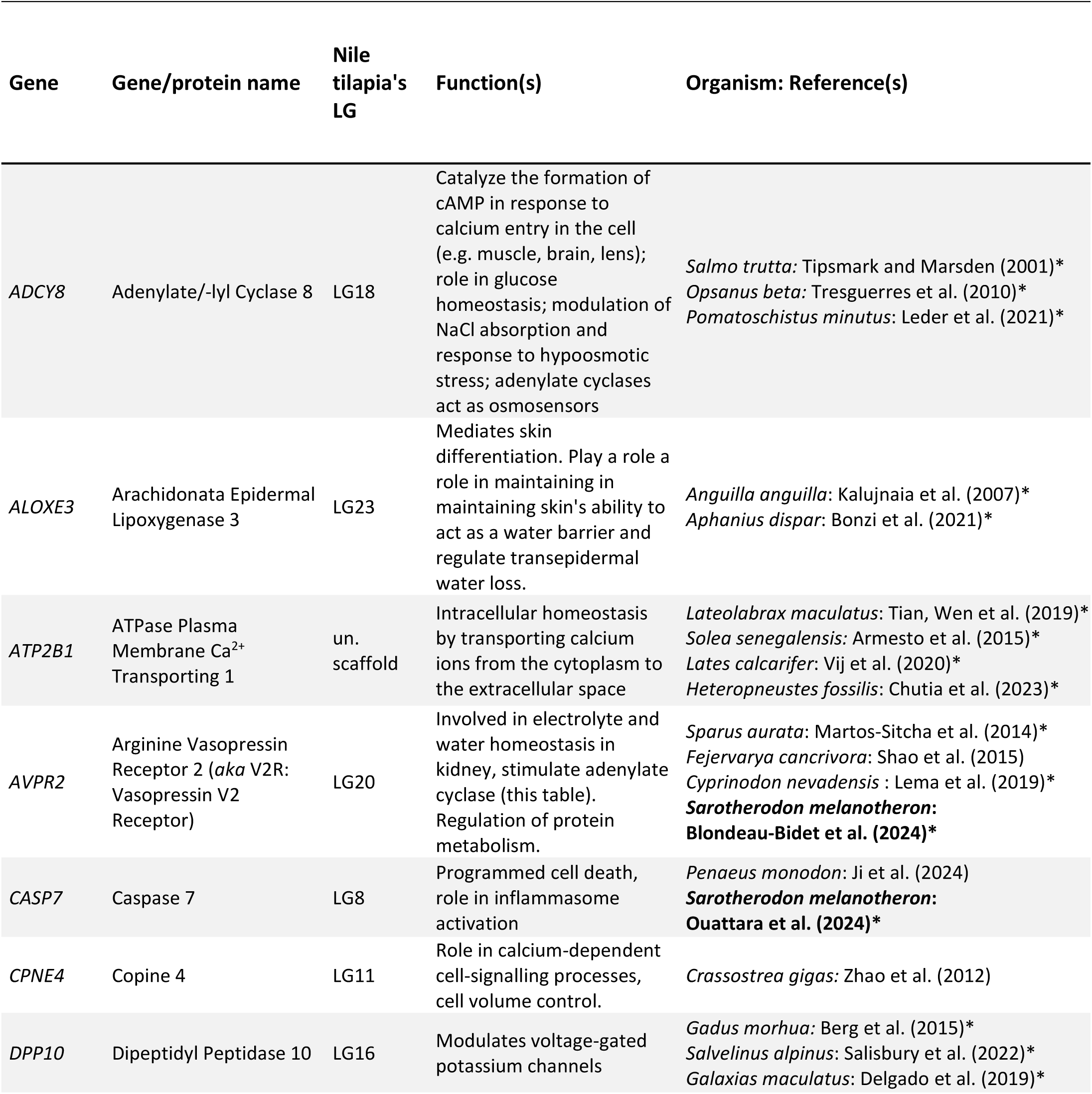
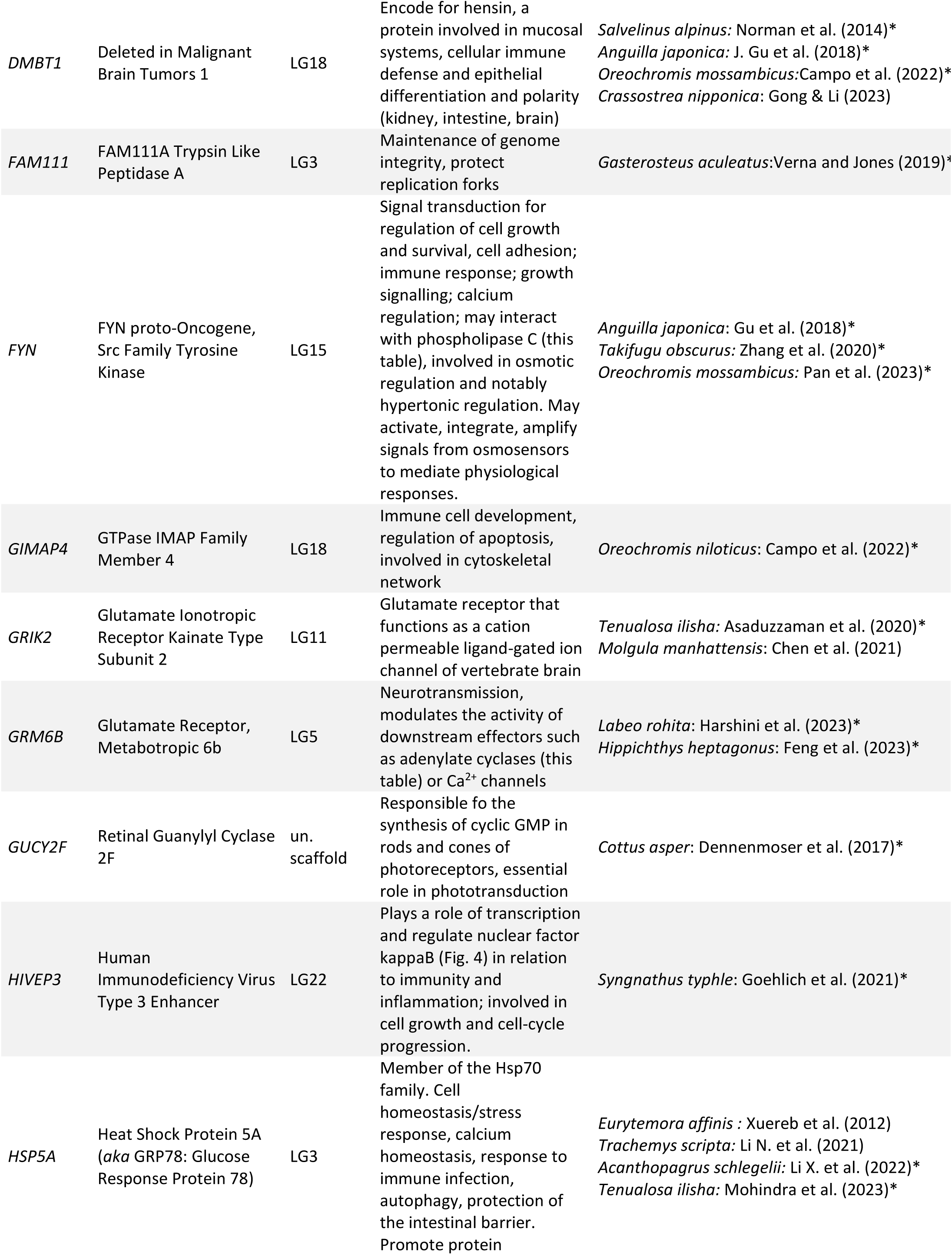
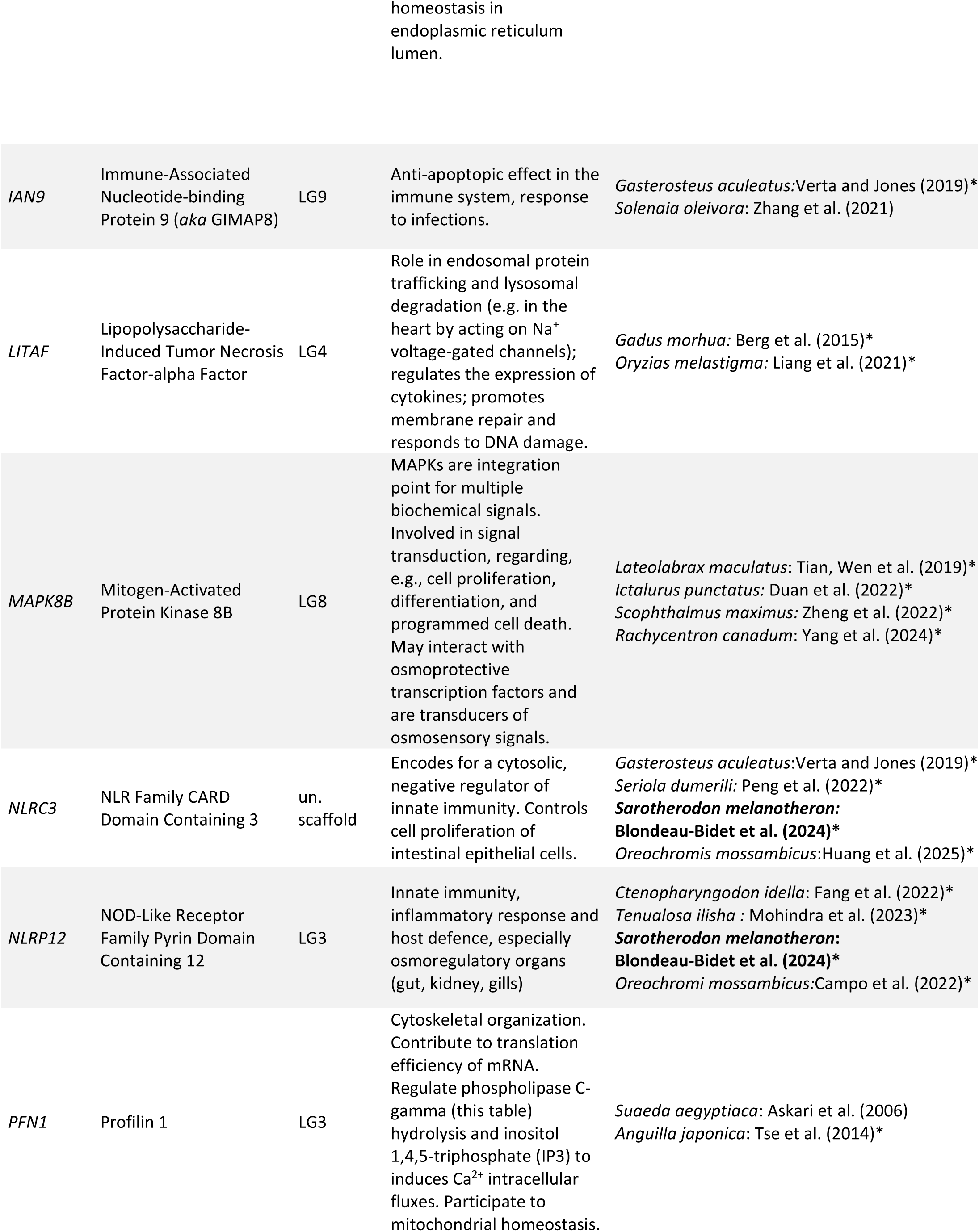
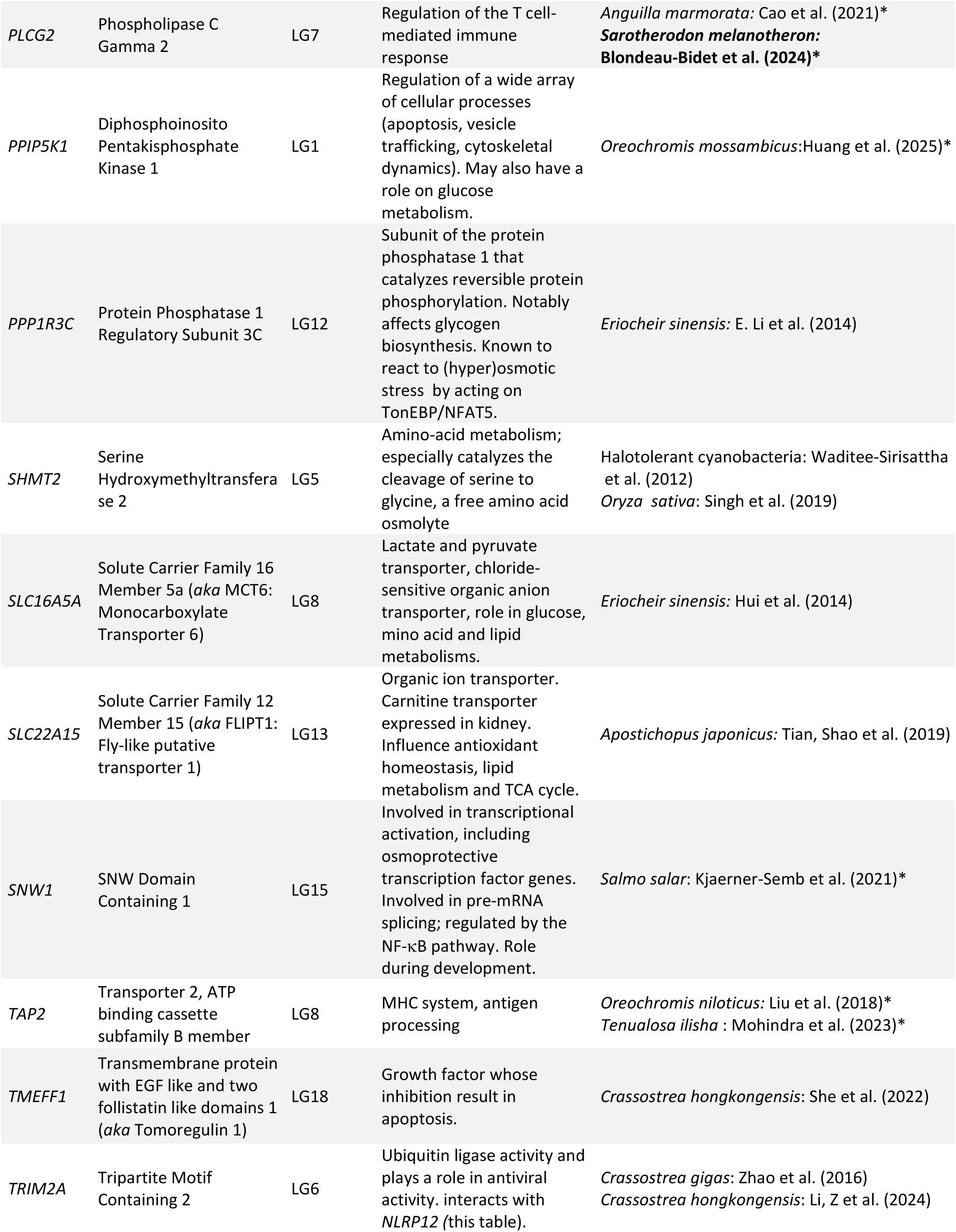
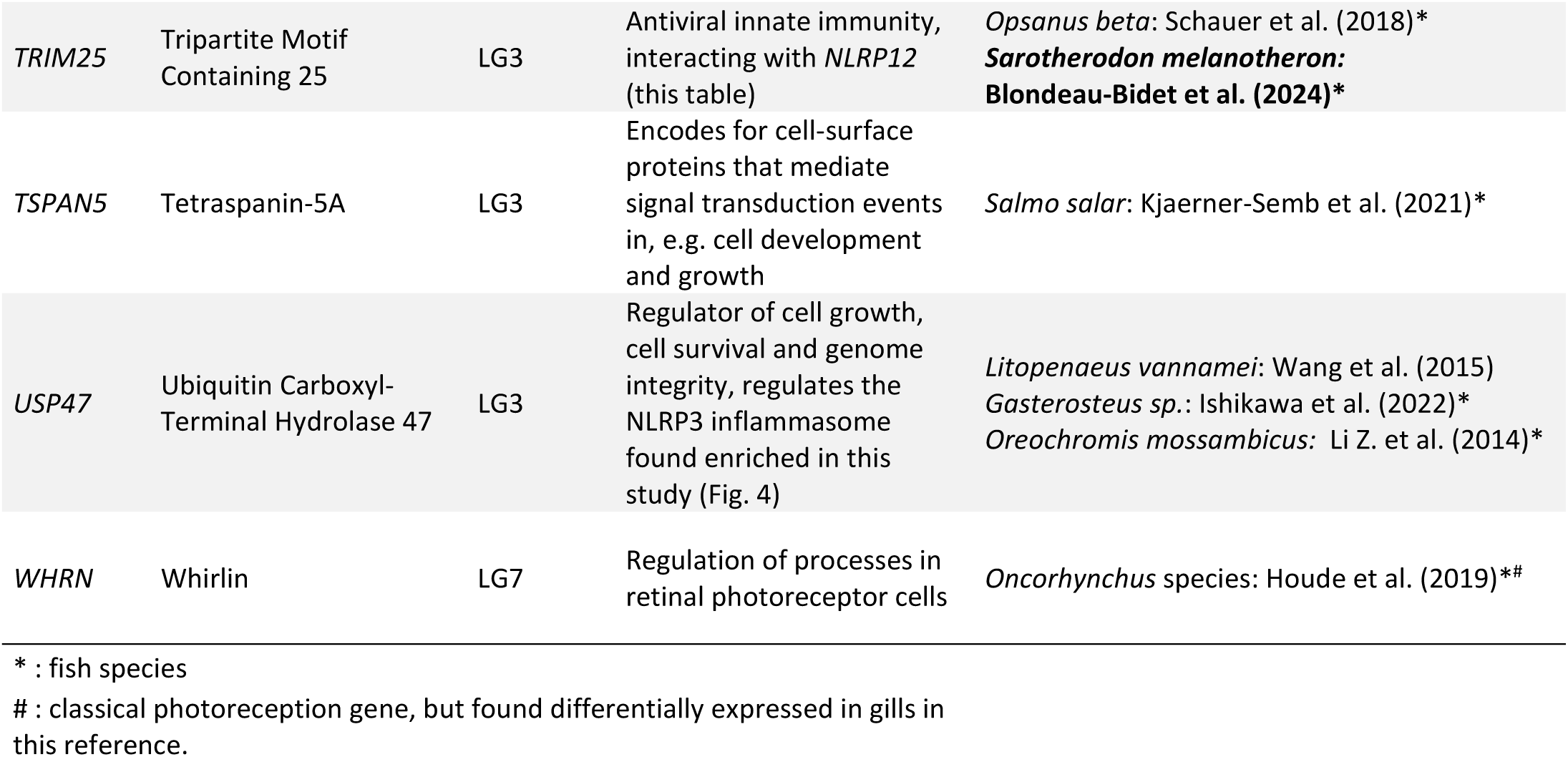
Alphabetical list of genes with at least one outlier SNP previously detected in gene expression or genomic studies interested in salinity response. The linkage group or unassembled scaffolds where these genes are located along the *O. niloticus* reference genome are given. Species names are given. Genes shown to be differentially expressed along the freshwater-hypersaline gradient in black-chinned tilapia are shown in bold. The full list of high *F*_ST_ outlier SNPs detected in this study is given in Supplementary Table S2.

## 4. Discussion

Except for a few genetic studies conducted in soda lakes (i.e. referring to a different type of salinity; Kavembe et al., 2014; Ford et al., 2015), this is the first study to investigate genetic differentiation at a genome-wide level in a West African euryhaline fish species along a fresh-to-hypersaline sodium chloride water gradient. No similar studies have been conducted on any other hypersaline fish species. Studies using a high-throughput gene expression approach or advanced proteomics are also rare, despite extended salinity gradients being expected to increase in the near future and affect many other aquatic species. This study aimed to evaluate the adaptive component of genetic differentiation in black-chinned tilapia across this gradient by identifying SNPs with high *F*_ST_ values (i.e. putative adaptive outliers that are responsible for increased genetic divergence) and the functional pathways involved in responding to environmental salinity and other energy-demanding processes.

### 4.1. Population genetic structure

Previous genetic studies using allozymes, mitochondrial DNA and a few other gene markers have examined the entire distribution range of the black-chinned tilapia (Pouyaud & Agnèse, 1995; Adépo-Gourène et al., 1998; Falk et al., 1999; 2003). However, microsatellite loci analyses, which are considered to provide a more detailed picture of genetic population structure, have mainly focused on *S. m. melanotheron* in the Gulf of Guinea, where salinity levels are much lower than in the Saloum (Pouyaud et al., 1999; Agnèse et al., 2008; Yoboué et al., 2012; 2014; Amoussou et al., 2018). Only one study has considered Senegambian samples belonging to the *S. m. heudelotii* subspecies (Ndiaye, 2012), including the five Senegalese samples used in the present study (Fig. 1). As in other cichlid species (e.g. *Pelvicachromis taeniatus*: Langen et al., 2011; *P. pulcher*: Schons et al., 2022; *Neolamprologus pulcher*: Hellmann et al., 2016), as well as in previous studies of black-chinned tilapia (Pouyaud et al., 1999; Agnèse et al., 2008; Ndiaye, 2012; Amoussou et al., 2018), we observed departures from HWE in most, but not all, samples. Our observations are therefore consistent with previous findings in this species, and classical mechanisms have been proposed to explain these deviations. These include relatedness between close relatives (Pouyaud et al., 1999), selection for homozygotes (Yoboué et al., 2012), the sampling of partially genetically differentiated subpopulations (i.e. the Wahlund effect), and the presence of null alleles (Agnèse et al., 2008; Amoussou et al., 2018). However, the issue of HWE in *S. melanotheron* requires further research and is beyond the scope of this study.

We have demonstrated significant genetic differentiation across the genome among Senegambian samples of black-chinned tilapia, and our results corroborate patterns identified by Ndiaye (2012) using just six neutral microsatellite loci. Our results confirm the following: *(i)* drainage basins, such as Hann Bay, an isolated SW population, represent distinct genetic units in black-chinned tilapia; *(ii)* there is no isolation by distance, primarily because *(iii)* southern (Gambia Basin) and northern (Guiers Lake) samples cluster together, as demonstrated by the NJ tree and main PCA scatterplot (Figs. 2A and 3A). Pouyaud & Agnèse (1995) also observed the clustering of remote samples from the Gambia and Senegal basins using allozymes. In addition to geographical differentiation, the full set of SNPs revealed that the Saloum samples were distributed along PCA2, a salinity gradient axis (Fig. 3A). However, this axis does not strictly reflect the actual intensity of the hypersaline gradient observed in the estuary. Only the PCA restricted to high *F*_ST_ outlier loci suggested that the Saloum samples followed the observed hypersaline gradient. However, this pattern may more broadly reflect an ordination that separates samples exposed to constant FW or SW conditions from those experiencing salinity fluctuations (Fig. 3C). Results from the pairwise *F*_ST_, NJ tree, admixture analysis and PCA scatterplots suggest some gene flow within the Saloum, primarily between the Missirah and Foundiougne samples and, to a lesser extent, between the Kaolack hypersaline samples. While these methods vary in their spatial and temporal resolutions, the genomic patterns observed at outlier loci were consistent with the findings of Panfili et al. (2005) and Diouf et al. (2006). These authors reported reduced connectivity among the estuarine populations of black-chinned tilapia in the Saloum using, respectively, fluctuating asymmetry in otolith shape and otolith strontium-to-calcium concentration ratios.

Nevertheless, Lake Guiers is the most differentiated sample when outlier loci are used, suggesting that the effects of constant FW conditions may outweigh those of hypersalinity in our dataset. This implies that strong local adaptive patterns of salinity-driven genomic variation are reduced when gene flow is sufficient within a single, open, wide estuary system, such as the Saloum. The clearest examples of local adaptation in fish are generally found in landlocked populations where connectivity between saline and non-saline environments has been disrupted (e.g. salmonids: Kjaerner-Semb et al., 2021; Salisbury et al., 2022; galaxids: Tuiz-Jarabo et al., 2016; mugilids: Blel et al., 2010; clupeids: Han et al., 2025). This is also the case for black-chinned tilapia in the Ivory Coast (Agnèse et al., 2008), and the Guiers Lake sample in our study may also illustrate this general observation. Black-chinned tilapia from this lake have had to adapt to constant FW conditions since the completion of the Diama dam on the Senegal River in 1985 (Cogels et al., 1993). Prior to the construction of the dam, the Senegal River and Lake Guiers were affected by seawater and the salinity could fluctuate seasonally from fresh to brackish water (15–20 ‰) (Gac et al., 1986). These results suggest a relatively rapid adaptive response in Lake Guiers, while the background genome remains largely consistent with that of most other samples along PCA1. These samples account for the majority of the observed genomic variation among black-chinned tilapia (Fig. 3A).

As observed in other coastal and estuarine organisms (e.g., Beheregaray & Sunnucks, 2001; Nielsen et al., 2020; Rautsaw et al., 2021; Chen et al., 2023; dos Santos Longo et al., 2024), our study highlights the difficulty of distinguishing between the environmental and geographic factors responsible for genetic differentiation in black-chinned tilapia. However, it refutes the hypothesis that extensive phenotypic plasticity alone accounts for the observed trait variation among Senegambian populations along the salinity gradient. Although definitive evidence of local adaptation to hypersaline conditions is lacking, our findings call into question the idea that climate change has driven a genomic response specifically linked to the hypersaline gradient that has developed in the Saloum estuary over the last fifty years. At this stage, it seems reasonable to interpret the observed adaptive genomic variation as occurring across the fresh-to-hypersaline gradient, rather than being specific to hypersaline environments. Further investigation is required, including temporal replication of this study by comparing 2006 samples with contemporary ones (i.e. spanning 60–80 generations). Additional insights could be gained through common garden experiments (a review in Oomen & Hutchings, 2015) or reciprocal transplants (Brennan et al., 2015).

### 4.2. Support for salinity-based adaptive variation

Although most outlier SNPs did not map to annotated genes in the Nile tilapia reference genome, our RADseq data revealed candidate genes and pathways that are potentially involved in salt stress responses. Many of these are also known to be involved in salt stress responses in other aquatic organisms, including *S. melanotheron* (Table 2). We present selected examples in Fig. 4 and Table 2, acknowledging that other relevant genes, some of which are also involved in salt stress in plants, were identified. While overlaps between gene expression and outlier loci are generally mutually exclusive in fish (e.g. Kozak et al., 2014; Thorstensen et al. 2021), our results suggest some alignment between the genomic and transcriptomic signals.

We found significant enrichment of pathways involved in the dynamic assembly and disassembly of protein complexes that mediate most intracellular signalling pathways and signal transduction. These pathways are key to the ability of cells to survive and respond to a changing environment. These findings suggest links to osmosensing (Kültz, 2012) and protein quality control under osmotic stress (Tang & Lee, 2013; Tsai et al., 2018). The GO term associated with “protein modification by conjugation or removal” refers to post-translational protein modification, which is critical for responding to salinity fluctuations (Leprêtre et al., 2025). This includes, among other processes, ubiquitination, and this GO term has been found to encompass four genes that encode E3 ubiquitin proteins (HACE1, RNF19B, TRIM2 and TRIM25; Table 2 and Suppl. Mat. Table S2). These proteins are the final enzymes that regulate the degradation of proteins, thereby preventing their accumulation in epithelial cells and also controlling cell growth and apoptosis in response to environmental changes (Sardella & Kültz, 2009; Tang & Lee, 2013; Tsai et al., 2018). Using Metascape, we identified the most relevant enriched term as ‘regulation of protein-containing complex assembly’, which is also associated with ‘endocytic recycling’ (Suppl. Mat. Table S3). This association has been repeatedly observed in the context of salinity variation and adaptive euryhalinity in fish (e.g., Tine et al., 2014; Cozzi et al., 2015; Li & Kültz, 2020), as well as to specific salinity adaptation in hypo-*or* hyperosmotic environments (Guo et al., 2018; De Vos et al., 2021; Zong et al., 2021).

We also identified the inositol pathway, which is known to be central to osmoregulatory responses in fish at the cellular level (Cui et al., 2022), including in response to hypersalinity (e.g., Fiess et al., 2007; Dowd et al., 2010; Sacchi et al., 2013; Bu et al., 2021). This pathway regulates cell signalling and energy homeostasis and coordinates metabolic processes (Shears, 2018), including carbohydrate metabolism in fish subjected to salt stress (Zhu et al., 2021, 2022; Pan et al., 2023; F. Zhang et al., 2024). The inositol and phospholipase pathways, one of which was also found to be enriched in this study, interfere with each other in mammals (Kim et al., 2024) and with components of the immune response. The phospholipase pathway has also been shown to be affected by salinity gradients (Cao et al., 2021 ; Han et al., 2025), and the *PLCG2* gene associated with this pathway was found to be differentially expressed in the gills of black-chinned tilapia (Blondeau-Bidet et al., 2024). Phospholipase C genes regulate e.g. calcium storage and release or are regulated (e.g. by prolactin) in tilapias (Tipsmark et al., 2005). Another outlier gene detected in this study is *PPIP5K1*, which belongs to the inositol pathway and is involved in Na^+^/K^+^-ATPase degradation (Table 2). It also interferes with cytokines in immunoregulation (Pulloor et al., 2014), lipid metabolism and cytoskeletal organisation (Machkalyan et al., 2016), all of which have been found to be differentially expressed in the gills of *S. melanotheron* (Blondeau-Bidet et al., 2024). At the protein level, PPIP5K1 interacts with SHMT2 (serine hydroxymethyltransferase 2), which is an outlier that was also identified in this study (Gu et al., 2021; Table 2). Mitochondrial SHMT2 catalyzes the interconversion of serine and glycine, with glycine — a free amino acid and osmolyte — being involved in the metabolic response to hyperosmotic stress (Martins et al., 2011; Glover et al., 2017). Further research is needed to investigate the relationship between SHMT2 and osmotic stress.

Among other pathways, the involvement of the thyroid axis (KEGG hsa04918; Fig. 4) in response to salt stress and actions on the Na^+^/K^+^-ATPase sodium pump are established (Klaren et al., 2007; Deal & Volkoff, 2020), and notably in the response of *O. mossambicus* to hypoosmotic conditions (Peter et al., 2000; Seale et al., 2021). Cell adhesion and synapse organization pathways found in this study are supported by the available literature on osmoregulation and osmosensing (Denker & Barber, 2002; Bourque, 2008, respectively) and found significantly enriched in black-chinned tilapia across the fresh-to-hypersaline water gradient (Blondeau-Bidet et al., 2024), but also in *O. mossambicus* to sense and respond to abiotic stimuli in both hypo- and hyperosmotic conditions (Seale et al., 2012; Gardell et al., 2013, respectively). Cell cycle-related pathways are crucial to the cellular stress response when facing salinity challenges (e.g., Evans & Kültz, 2020), as especially shown in *O. mossambicus* in hypersaline conditions (Kammerer et al., 2009).

As already suggested, the mapping of outlier SNPs to genes associated with immune pathways were also found in our study. Gene expression studies reporting co-variation in immune and osmoregulatory-related pathways are numerous in euryhaline fish within the FW-SW range, notably in tilapias (e.g., Campo et al., 2022; Farhadi et al., 2023; J. Zhang et al., 2024). This relationship between salinity and immune responses extends to hypersaline water >40 ‰ in fish (*G. aculeatus*: Li & Kültz, 2020; *F. heteroclitus*: Tao & Breves, 2024) or bivalves (*Crassostrea corteziensis*: Pérez-Velasco et al., 2022), and more generally in organisms living in extreme environments (Tong and Li, 2020; Lu et al., 2022). Genomic studies also reported SNPs in immune genes when targeting salinity-based adaptation (Gao et al., 2021a; Velotta et al., 2022). Our enrichment analysis is also consistent with the RNA sequencing study of *S. melanotheron* gill tissues, which reported enrichment of immune response-related functions (e.g., NF-κB signaling; Fig. 4) along the FW-hypersaline gradient (Blondeau-Bidet et al., 2024). This co-variation presumably occurs because immune responses involve trade-offs with other energetically costly physiological mechanisms during prolonged cellular stress (Lazzaro & Little, 2009), including the salinity stress response (Wilck et al., 2019). For example, the activation of the NLRP3 inflammasome (Fig. 4) relies on the ability to sense numerous structurally diverse stimuli (Chen & Nuñez, 2010) and requires coordinated changes of intracellular ion concentrations (e.g., Ca^2+^ mobilisation, K^+^ and Cl^-^ efflux; Gong et al., 2018; C. Li et al., 2021), which are likely to trade off with osmotic regulation. At the gene level, these include in particular *NLRP12* (Table 2) which belongs to a salt-sensitive NOD-like receptor gene family involved in immunity and acts as a negative regulator of the NF-κB signalling pathway (Chuphal et al., 2022; Wan et al., 2024). *NLRP12* is known to interact with the aforementioned *TRIM2* and *TRIM25* genes, but also with another NLR gene (*NLRC3*) (Morimoto et al., 2021; Liu et al., 2023) that was also found to contain one outlier locus in this study (Table 2). *NLRP12* and *NLRC3* were both found differentially expressed in *O. mossambicus* and *S. melanotheron* as a function of salinity (Table 2).

The enriched VEGFA-VEGFR2 pathway (vascular endothelial growth factor A [receptor 2]; Fig. 4) found in this study mediates cellular responses involved in angiogenesis and vascular permeability, and is affected by osmotic conditions (Manolopoulos et al., 2000; Qin et al., 2020). This suggests some functions related to vasoconstriction or vasodilation under salinity stress, whereas to our knowledge this pathway has never been studied in detail in fish and has only been reported anecdotally (Gao et al., 2021a; Pan et al., 2024). However, VEGFA has been shown to react to *myo*-inositol in gill and kidney of turbot (*Scophthalmus maximus*; Cui et al., 2020). This pathway may deserve more attention as it is also known to be regulated by prolactin and growth hormone (Clapp et al., 2009), which are differentially regulated along the salinity gradient in the black-chinned tilapia (Tine et al., 2007, 2012). Other significantly enriched pathways found in this study may also deserve increased attention in the future. For example, photoperception-related pathway (Fig. 4) has been shown to be relevant in the response to salinity in fish (e.g. *Coilia nasus*: Gao et al., 2021a, b), perhaps because of its dependence on prolactin production across the salinity gradient (Pavlova et al., 2022).

### 4.3. Remaining questions

Beyond the classical limitations of genome scan studies of adaptive variation using RADseq (e.g., detection of false positives, allele drop-out, sample sizes, underlying models; Gautier et al., 2013; Mastretta-Yannes et al., 2014; Lotterhos & Whitlock, 2015), our study based on SNPs with high *F*_ST_ estimates is biased towards detecting outlier SNPs under putative positive selection and neglects those under weaker selection pressure or balancing selection that may be important for driving salinity-based adaptation (Durland et al., 2021; Lee et al. 2022). Nevertheless, a PCA-based strategy favoring loci with high *F*_ST_ values has already been shown to be effective in salinity-based fish studies (*Clupea harengus*; Han et al., 2020). Furthermore, any relative weakness is offset by enrichment analysis, which identifies pathways and outlier-containing genes that have already been found to be involved in the salinity response. Lastly, well-documented and not only putative positive selection has already been reported in euryhaline fish for some of outlier-containing genes outlined in Table 2 (*ATP2B*: Velotta et al., 2022; *AVPR2*: Shao et al., 2015; *HIVEP3*: Diepeveen et al., 2013; *GRM* and *SLC22* gene families: Han et al., 2025; Zhou et al., 2023, respectively) and gives support to our results.

We have found that it is difficult to consistently distinguish the pathways or genes that contribute to the adaptation to the hypo-vs the hypersaline environment. For instance, the inositol-related pathways have been shown to respond predominantly to hyperosmotic environments (Li & Evans, 2020), while the thyroid hormone synthesis pathway responds to hypoosmotic environments (Seale et al., 2021 ; Han et al., 2025), but we do not have direct confirmation for *S. melanotheron*, as these pathways were not found to be down- or up-regulated along a 0-80 ‰ gradient (Blondeau-Bidet et al., 2024). Furthermore, LG18, but also - with lower effect sizes - LG4, LG11 and LG16, have been identified as physical locations for quantitative trait loci (QTL) involved in the response to salt stress in *Oreochromis spp* (X.H. Gu et al.,2018; Jiang et al., 2019; Yu et al., 2022; Huang et al., 2024; De Liu et al., 2025; Z. Yang et al., 2025). We found in this study that these QTLs are depleted for the outlier loci. Only one outlier SNP mapped to the QTL identified by Yang et al. (2025) on LG11. This SNP was associated with the *CPNE4* gene (Table 2) that encodes for a calcium-dependent membrane-binding protein related to the regulation of cell volume control, one important cellular component of osmotic regulation. This lack of overlap between studies remains intriguing, as patterns of intra- and inter-chromosomal variation are reduced in cichlids (Conte et al., 2019), as shown for LG1 of *O. niloticus* and *S. melanotheron* by Gammerdinger et al. (2016). It could be hypothesized that our study covered a much wider range of salinities than the QTL studies in *Oreochromis spp*. that never considered experiments >42 ‰. Other LGs might be involved at higher salinities, depending on the genomic architecture of the species that is responsible for the observed trait variation. Yu et al. (2022) effectively reported a salinity-based QTL on LG18 in a tilapia cross and showed that this QTL shifted around 13-14 Mb vs 19-27 Mb in other studies, i.e., in a genomic region in which we detected one outlier SNP in the *DMBT1* gene whose expression has been found associated with salinity in *O. mossambicus* (Campo et al., 2022), and other species (Table 2). *DMBT1* encodes for hensin, an extracellular matrix protein involved in cell polarity, in epithelial intercalated cells of the kidney where it participates to the regulation of water and acid-base balance (Gao et al., 2010). *DMBT1* is then relevant to adaptively respond to osmotic modification. Overall, a better understanding of the black-chinned tilapia’s specific response to salinity variation requires improved knowledge of its genome. This would provide valuable insight into the mechanisms of physiological adaptation, as has been achieved in other estuarine species (*F. heteroclitus*; Reid et al., 2017). QTL mapping and genome resequencing approaches (Laporte et al., 2015; Brennan et al., 2018) could also be promoted in *S. melanotheron*.

## 5. Conclusion

An adaptive component of genome-wide genetic differentiation was detected in the black-chinned tilapia across the Senegambian fresh-to-hypersaline gradient, but the adaptive genomic variation is not specific to hypersaline environments. Several outlier loci mapped to genes linked to salinity adaptation, partially consistent with previous high-throughput RNA sequencing data established along this gradient in this species. However, while RADseq reveals key signals, it remains insufficient to fully resolve the genomic basis of evolutionary mechanisms underpinning extreme euryhalinity in *S. melanotheron*. The role of rapid, hypersalinity-driven adaptation due to climate change remains plausible but requires further investigation. Projected increases in estuarine salinity (Bodian et al., 2018; Trisos et al., 2022) highlight the need for deeper insights into adaptive responses, which could inform breeding programs for salinity-tolerant strains, essential for food security in Africa and beyond (Yue et al., 2024).

## Acknowledgments

The MGX facility (Montpellier, France) participated to Illumina sequencing and acknowledges financial support from the France Génomique national infrastructure, funded as part of the Agence Nationale de la Recherche “Investissement d’Avenir” (ANR-10-INBS-09). Authors thank E. Blondeau-Bidet, S. Guendouz and M. Simier for support and help at various steps of this work.

## Data Availability

The molecular data that support the findings of this study are openly available in NCBI (https://www.ncbi.nlm.nih.gov/) under accession PRJNA1143382.

## Captions of Supplementary Materials

**Supplementary Figure S1:** Distribution of FST values for the 16,786 SNPs analyzed in this study.

**Supplementary Table S1:** Description of individual samples used in this study including the number of sequencing reads and individual call rates. Individuals that have been discarded are reported in red.

**Supplementary Table S2:** Distribution of outlier SNPs and position along chromosomes (linkage groups: LG) and unassembled scaffolds of the Nile Tilapia (Oreochromis niloticus, accession number: GCF_001858045.3) based on PCA. Gene ID is reported and the nature of the region each SNP mapped is reported when available (gene, protein coding region, pseudogene, lncRNA, miRNA…). When one outlier SNP mapped to an annotated gene, the gene symbol (from www.genecards.org) and the gene name are given. Genes that were not consistently annotated are reported in italics.

**Supplementary Table S3:** Output of the Metascape analysis based on annotated genes containing retained outlier SNPs. Cells in red in the ȘEnrichmentș spreadsheet correspond to the top 20 GO terms (Fig. 4 of the manuscript).

