## Supplementary figures and images for "Genomic differentiation of the black-chinned tilapia (*Sarotherodon melanotheron*) along a fresh-to-hypersaline water gradient"

### Supplemental Figure S1

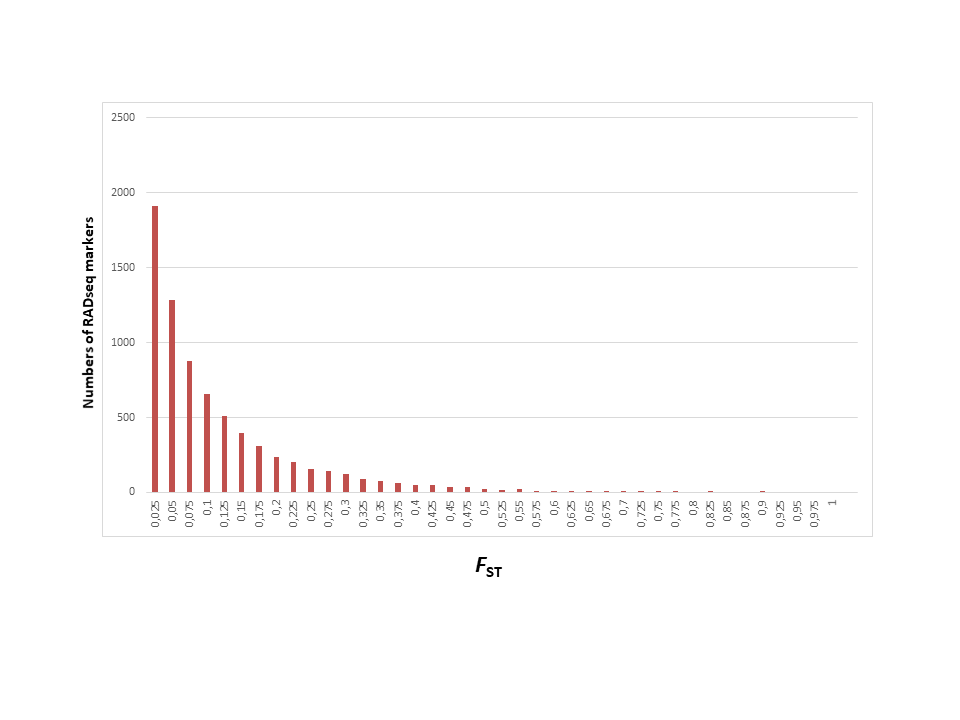
